# Biochemical reactions between intra-articular tissues and joint instability in a rat model of osteoarthritis

**DOI:** 10.1101/2023.03.28.533768

**Authors:** Kenji Murata, Sora Kawabata, Takuma Kojima, Yuichiro Oka, Chiharu Takasu, Hidenobu Terada, Naohiko Kanemura

## Abstract

**Aims:** Joint instability is associated with various joint conditions including osteoarthritis (OA) and inflammation, and we have developed model which is determined to role of knee instability. Investigating cartilage maintenance factors such as hyaluronic acid (HA) and glycosaminoglycans (GAGs) can provide insights into the effect of the mechanical stress and the inhibitor used, with the following aims: 1) whether cartilage degeneration is inhibited in the new model, 2) whether combination TGF-β1 inhibition mitigates cartilage degeneration, and to determine the role of TGF-β1 in synovitis using fibroblasts.

**Main methods:** We used this novel model to investigate inhibition of OA progression with a focus on HA and GAGs, which help maintain the cartilage and synovial membrane. In detail, mechanical tests, X-ray, histological, and protein and mRNA expression analyses were used to determine the role of joint stability using in vivo model or fibroblast from synovial membrane.

**Key findings:** Joint stability mitigated cartilage degeneration loss, decreased osteophytes, increased the expression levels of HA and GAGs in the synovial membrane, and decreased the release of pro-inflammatory factors in rats. Moreover, injection of TGF-β1 inhibitor in an inflammatory synovial membrane promoted HA and GAGs expression. In synovial fibroblast cells, inhibition of TGF-β1 over expression significantly inhibited the downregulation of pro-inflammatory factors and promoted the upregulation of lubrification for cartilage.

**Significance:** Our results suggest that joint instability is an independent mechanical factor for OA progression. The results provide novel insights into the association between OA and joint instability, which has significant human sciences implications.

**Research Highlights:** • Established a new experimental rat model of the different joint instability for elucidate osteoarthritis onset/progression

• Using Histological staining to investigated the osteoarthritis including synovitis and osteophytes of the novel model

• Using fibroblast from synovial membrane to investigated the fibrosis

• Joint instability exacerbates articular cartilage degeneration and decreases HA and GAGs protein expressions in synovial membrane

• TGF-β1 inhibitor on early osteoarthritis joints may suppress synovial inflammation

## 1. Introduction

Knee osteoarthritis (OA) interferes with daily living activities due to dysfunction and pain. The main pathology of OA involves cartilage degeneration, osteophytes, and synovitis, and has recently been reported as a continuous inflammatory disease in the knee joint[1–3]. Along with aging, sex[4], obesity[5,6], traumatic injury[7], and gene profiles[8], mechanical stress is an important risk factor for OA[9,10]. For example, excessive exercise loading increases mechanical stress, or lowering limb suspension decreases it for articular cartilage[11]. Mechanical stress models generated via surgery such as anterior cruciate ligament transection (ACL-T) and destabilization of the medial meniscus (DMM) have been used to study the progression to articular cartilage degeneration and synovitis[12,13].

Injuries and diseases associated with joint instability increase mechanical stress of knee joint, which is highly associated with OA. However, the role of joint instability during mechanical stress in OA has been previously investigated[14–17]. Joint instability, also referred to as joint laxity or hyper mobility, is an important factor in the treatment and management of joint conditions, and causes structural damage to the knee joint[18–20]. Soft or hard knee bracing can provide external stability and improve patients’ quality of life. Also, physical therapy may restore joint stability by strengthening articular muscles and increasing proprioceptive awareness[18,21,22]. However, the pathological influence of instability on articular cartilage or synovitis remains poorly understood. Therefore, understanding the pathology of joint instability-induced knee OA is essential. ACL transection in animal models has been widely used to investigate OA pathology[12,23,24]. However, the role of instability remains unclear. We have previously reported anterior cruciate ligament (ACL) healing post traumatic injury and developed a knee instability animal model. In this study, we used this model to investigate OA progression[25–27]. Since this model provides external stability, joint instability by ACL injury can be controlled. Our technique can be used to determine whether joint instability is an independent factor for OA.

Among the various factors that influence OA progression, transforming growth factor-β (TGF-β) plays an important role in both the cartilage and synovium[28–32]. Hyaluronic acid (HA) is important factor for homeostasis in articular cartilage, and has the simplest structure of all glycosaminoglycan (GAGs), which is important for lubricating joints. TGF-β1 promotes collagen type 2 or aggrecan synthesis, which is important when assessing cartilage histology[33]. Meanwhile, an over expression in TGF-β1 in the synovium membrane induces macrophages and inflammatory cytokines, thereby negatively affecting synovial fibrosis and inducing the formation of marginal osteophytes[34,35]. Controlling TGF-β1 may prevent intra-articular OA. Therefore, this study investigates the effect of OA on joint instability using a novel model, with the following aims: 1) to determine whether cartilage degeneration is inhibited in our model in which joint instability is controlled, 2) to determine whether TGF-β1 inhibition mitigates cartilage degeneration, and 3) to determine the role of TGF-β1 in synovitis using inflammation fibroblasts. Investigating cartilage maintenance factors such as HA and GAGs can provide insights into the effect of the mechanical stress and the inhibitor used in this study.

## 2. Material and Methods

### 2.1 Experimental design (in vivo study)

Male Wistar rats (10-week-old males; n = 56), weighting 220–250 g (Sankyo Labo Service Corporation INC., JPN), were randomly classified into 4 experimental groups, underwent ACL transection surgery and two surgical procedures to create different instability models. The animals were housed in a laboratory facility at 23 ℃ and 55% humidity in a 3-cage environment with free access to water and food. All animal protocols were conducted in accordance with the ARRIVE guidelines[36] and approved by the Ethics Committee of Saitama Prefectural University (approval number: 29-3). According to previously reported methods[37], 28 rats showed restoration of the knee joint kinematics, and the anterior instability of the tibia was limited following ACL surgical transection (CATI model). Meanwhile, the remaining 28 rats underwent sham ACL transection surgery (ACL-T model). Subsequently, both models were divided into different groups that either received a TGF-β1 receptor kinase inhibitor injection (SB-431542, Cayman Chemical, USA) or SHAM PBS. The rats were randomly assigned into four groups: CATI, CATI with inhibitor, ACL-T, ACL-T with inhibitor (n = 12 per group) (**Fig. 1A**), and were sacrificed 4 and 8 weeks post-surgery; the right knee joints of all rats were harvested for analysis.

**Fig. 1.**
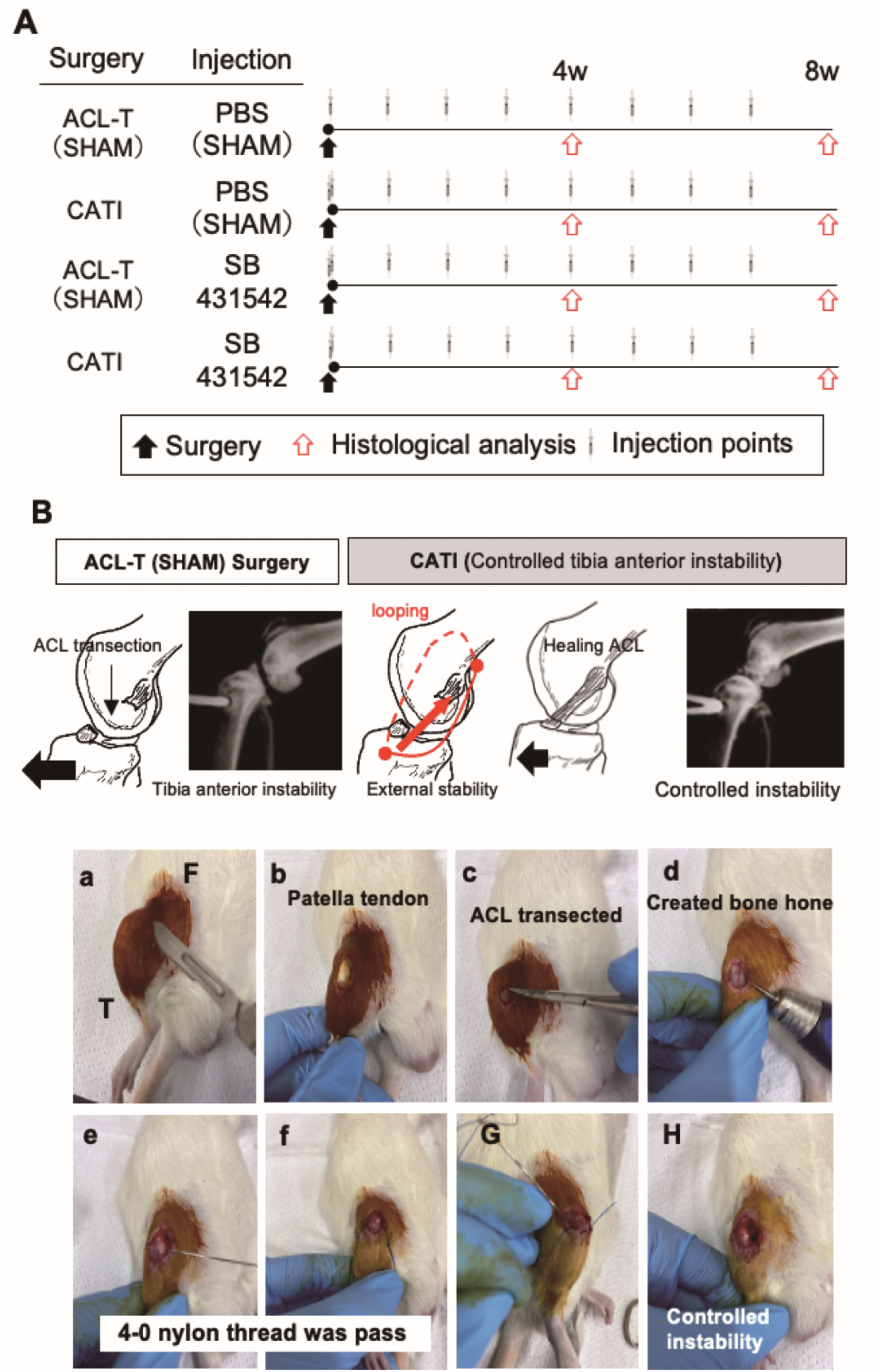
Study Design and surgery operation. (A) Schematic of the experimental design. ACL-T and CATI-operated rat were injected with the indicated concentrations of TGF-β1 receptor inhibitor times per week and sacrificed at 4 and 8 weeks after surgery (n = 14 rat per group). (b) Schematic of the anterior cruciate ligament transection (ACL-T) surgery model and controlled tibia anterior instability (CATI) surgery model. The CATI group involved a second procedure that was performed to restore biomechanical function following ACL transection using a nylon suture placed on the outer aspect of the knee joint. Under a combination anesthetic (medetomidine, 0.375 mg/kg; midazolam, 2.0 mg/kg; and butorphanol, 2.5 mg/kg), the medial capsule of the right knee joint was exposed without disruption of the patellar tendon (a,b), and the ACL was completely transected in both groups (c). After anterior tibial joint instability was confirmed, both groups underwent the creation of a bone tunnel along the anterior aspect of the proximal tibia (d), through which a 4-0 nylon thread was passed and tied to the posterior aspect of the distal femur(e,f). In the CAM group, the thread was tied tightly (g), dampening with abnormal joint movement without intra-articular suturing of the ligament (as performed in ACL reconstruction). In contrast, the nylon thread was not tightly tied in the SHAM group, and the anterior tibial joint instability remained. F, femur; T, tibia

### 2.2. Surgical procedures

According to previous studies[37], surgical procedures were performed under a combination anesthesia of 0.3 mg/kg medetomidine, 4.0 mg/kg midazolam, and 5.0 mg/kg butorphanol. Using a scalpel, a longitudinal incision on the internal margin of the patella was made to expose the patella tendon and synovial membrane, with ACL-T. All groups showed joint instability, then rat were undergo external stability surgery or sham surgery. A bone hole was transversely made on the tibia using a special machine, and controlled joint instability was achieved by passing a nylon thread in a loop between tibia and femur. The sham groups underwent the same procedure. Then, the surgical wounds were sutured using a silk suture, and the rats were allowed to fully recover (**Fig. 1B**).

### 2.3. Injection procedures

Rats were randomly treated with either the TGF-β1 receptor kinase inhibitor injection or vehicle. The treatment group received intra-knee injections of 25 μg/mL TGF-β1 receptor kinase inhibitor in DMSO (FUJIFILM Wako Pure Chemical CO., JPN) solution once a week.

### 2.4 Biomechanical test

Range of motion (ROM) and tibia anterior instability was analyzed using an X-ray machine (Softex INC., USA). The range of motion of the knee was measured at maximum flexion and extension following surgery, and calculated from the femur and tibia position using software (Image J Ver.1.52, JPN). The tibia anterior instability was measured in an anterior traction state using 0.2 N qualitative spring original machine at weeks 4 and 8, and calculated as previously described [37].

### 2.5 Histological analysis

Knee joint tissue samples were collected at 4 and 8 weeks; the tibia and femur were evaluated using Hematoxylin and Eosin (HE) staining for histological observation. The tibia was fixed in 4% paraformaldehyde (FUJIFILM Wako Pure Chemical CO., JPN) for 24 h, decalcified in 10% ethylenediaminetetraacetic acid (Merck, GER) solution for 60 days, and finally placed in the OCT compound to obtain frozen sections. Cartilage degeneration was evaluated using the Osteoarthritis Research Society International (OARSI) scoring system[38] in the medial central part of the articular cartilage and the posterior medial marginal region of the tibia and femur. Two authors (YO and TK) blindly evaluated all cartilage samples. The cartilage was evaluated at three sites (anterior and posterior, ×200 using a microscope), and three sections were used per knee.

### 2.6 Immunohistochemical examination

Sections were washed twice (5 min per wash) with phosphate-buffered saline (PBS), blocked with normal serum for 1 h at 22–24°C and incubated overnight at 4°C with rabbit polyclonal anti-Lubricin antibody (1:500, MABT401, MERCK, DEU). The sections were then washed three times with PBS, incubated with secondary antibody Dylight 649 mouse anti-goat (1:2000, Thermo Fisher Scientific, JPN) for 1 h at approximately 22–24°C and mounted with 4′, 6-diamidino-2- phenylindole (DAPI) (Vector, Burlingame, USA). Samples were observed under a BZ-X700 microscope (Keyence, JPN). Lubricin density in the cartilage was quantified using Image J software, while synovial membrane staining was semi-quantified. The results were blindly obtained and evaluated by one of the study authors.

### 2.7 Enzyme-linked immunosorbent assay (ELISA) for synovial membrane

Synovial membrane was collected at 4 and 8 weeks, and the levels of HA, Interleukin (IL)-1β, tumor necrosis factor (TNF)-α, TGF-β1, and collagen types 1 and 3 in the synovial membrane were measured using ELISA. Total protein samples (n = 5 per group) were prepared by homogenizing the synovial membrane tissue, excluding the meniscus and bone tissue, in a tissue protein extraction reagent (T-PER, Thermo Fisher Scientific, JPN) containing a protease inhibitor cocktail (Halt Protease Inhibitor Cocktail, Thermo Fisher Scientific, JPN). The protein concentration was adjusted using BCA method. The protein levels of TGF-β (88-50680-22, Thermo Fisher Scientific, JPN), HA (Hyaluronan Quantikine ELISA Kit, R&D systems, USA), IL-1β (88-6010-22, Thermo Fisher Scientific, JPN), TNF-α (88-7340-22, Thermo Fisher Scientific, JPN), collagen type 3 (MBS2509319, MyBioSource, USA), and collagen type 1 (MBS262647, MyBioSource, USA) were evaluated using ELISA kits in accordance with the manufacturer’s instructions.

### 2.8 Biochemical assays to measure GAG expressed of synovial membrane

Sulfated glycosaminoglycans (S-GAG) in medium was measured using Blyscan™ Glycosaminoglycan Assay Kit according to the manufacturer instructions (B1000, Biocolor Ltd., USA).

### 2.9 Extraction of rat fibroblast from synovial membrane (in vitro)

Fibroblast were harvested from the knee synovial membrane of male Wistar rats (n=8, aged 8 weeks). Synovial membranes were rinsed with Dalbeco-phosphate-buffered saline (D-PBS, Thermo Fisher Scientific, JPN) and minced with MEM-α containing 3 mg/mL collagenase V (FUJIFILM Wako Pure Chemical CO., JPN) on a shaker at 37 °C for 3 h. The solution was centrifuged at 300 ×*g* for 3 min to remove the supernatant, which was mixed with 1 mL α-MEM (including antibiotics and 10% FBS and Penicillin-Streptomycin) solution and seeded in 6 well dishes under a humidified atmosphere of 5% CO_2_ at 37 ℃. The second passage cells were used in the experiment.

### 2.10 Fibroblast treatment using IL-1β, TGF-β1, and SB-431542

Cultures were stimulated with/without recombinant rat IL-1β (1 ng/mL) (FUJIFILM Wako Pure Chemical CO., JPN), recombinant human TGF-β1 (10 ng/mL) (Peprotech, USA). Moreover, SB-431542 (10 μM) (Cayman Chemical, USA) was used to selectively inhibit TGF-β1/ALK5-smad2/3 signaling cascade. The second passage fibroblast were previous seeded 1×10^5^/mL into a 6 well plate in a 5% CO_2_ at 37 ℃ incubator, with 70–80% confluency.

### 2.11 Cell proliferation assay

Fibroblast were seeded (100 µL; 1.0×10^5^/mL) in Nunc 96-well plates (167008, Thermo Fisher Scientific, JPN) overnight under a humidified atmosphere of 5% CO_2_ at 37 ℃. After 12 h, the medium was treated with TGF-β1 and SB-431542 and incubated for 24 h. On the following day, 10 µL Cell Counting Kit-8 (CCK8, Dojindo Inc., JPN) reagents were added to each well and incubated for an additional 1 h. The absorbance was measured at the wavelength of 450 nm using plate reader.

### 2.12 Real-time reverse transcription polymerase chain reaction

The mRNA expression levels from the synovial fibroblasts were evaluated with a real-time reverse transcription-polymerase chain reaction (qPCR). Total RNA of cultured cells was extracted using RNeasy mini kit (QIAGEN, JPN) according to manufacturer’s instruction. The cDNA was synthesized using the High-Capacity cDNA Reverse Transcription Kit (4374966, Thermo Fisher Scientific, JPN). The taqman gene expression assay was used to amplify the target genes (*Il6*: Rn01410330_m1, *Prg4*: Rn01490812_m1, *Has1*: Rn01455687_g1, *Has2*: Rn00565774_m1, *Col1a1*: Rn01463848_m1, *Col3a1*: Rn01437681_m1). The gene expression was quantified by normalizing to the *β-actin* gene expression using the optimized comparative cycle threshold.

### 2.13 Biochemical assays to measure GAG expressed in treatment medium

S-GAG in medium was measured using Blyscan™ Glycosaminoglycan Assay Kit (B1000, Biocolor Ltd., USA). The media were centrifugation at 13,000 ×g to remove the pellet, and the supernatant medium was collected. A total of 100 µL medium was used for analysis from each culture, and the assessment was performed as in the in vivo study.

### 2.14 ELISA of fibroblast

Proteins in each synovial membrane fibroblast were harvested in a RIPA lysis buffer (89900, Thermo Fisher Scientific, JPN). The concentration was evaluated using Pierce™ Rapid Gold BCA Protein Assay Kit (A53226, Thermo Fisher Scientific, JPN). The protein concentration was adjusted using the BCA method. The protein levels of TGF-β (88-50680-22, Thermo Fisher Scientific, JPN), HA (Hyaluronan Quantikine ELISA Kit, R&D systems, USA), IL-1β (88-6010-22, Thermo Fisher Scientific, JPN), TNF-α (88-7340-22, Thermo Fisher Scientific, JPN), collagen type Ⅲ (MBS2509319, MyBioSource, USA), and collagen type Ⅰ (MBS262647, MyBioSource, USA) were evaluated using ELISA kits in accordance with the manufacturers’ instructions.

### 2.15 Statistical analysis

Data were analyzed using IBM SPSS Statistics 25 (IBM Japan, JPN), and tested for normality using the Shapiro-Wilk test. For the parametric data, one-way or two-way analysis of variance (ANOVA) was used in each group at different time points for each parameter. Nonparametric data were used to determine the effects of each group for each parameter using the Kruskal-Wallis test. The post-hoc test used was either the Tukey test for parametric data or Mann-Whitney U test with Bonferroni-Holm for non-parametric data correction according to the homogeneity test. Parametric data are presented as mean (95% confidence interval: CI), while nonparametric data are presented as median with interquartile ranges [IQR]. *p* < 0.05 were considered statistically significant.

## 3. Results

### 3.1. Animal condition

No significant changes in body weight before and after surgery, or surgery and injection at 4 (*p* = 0.100) and 8 (*p* = 0.330) weeks were observed (**Fig. 2A**). Postoperative urinary C-reactive protein was high in all groups on day 4, yet no differences between the groups were observed (4 day: *p* = 0.349, week 2: *p* = 0.100, and week 4: *p* = 0.330; **Fig. 2B**)

**Fig. 2.**
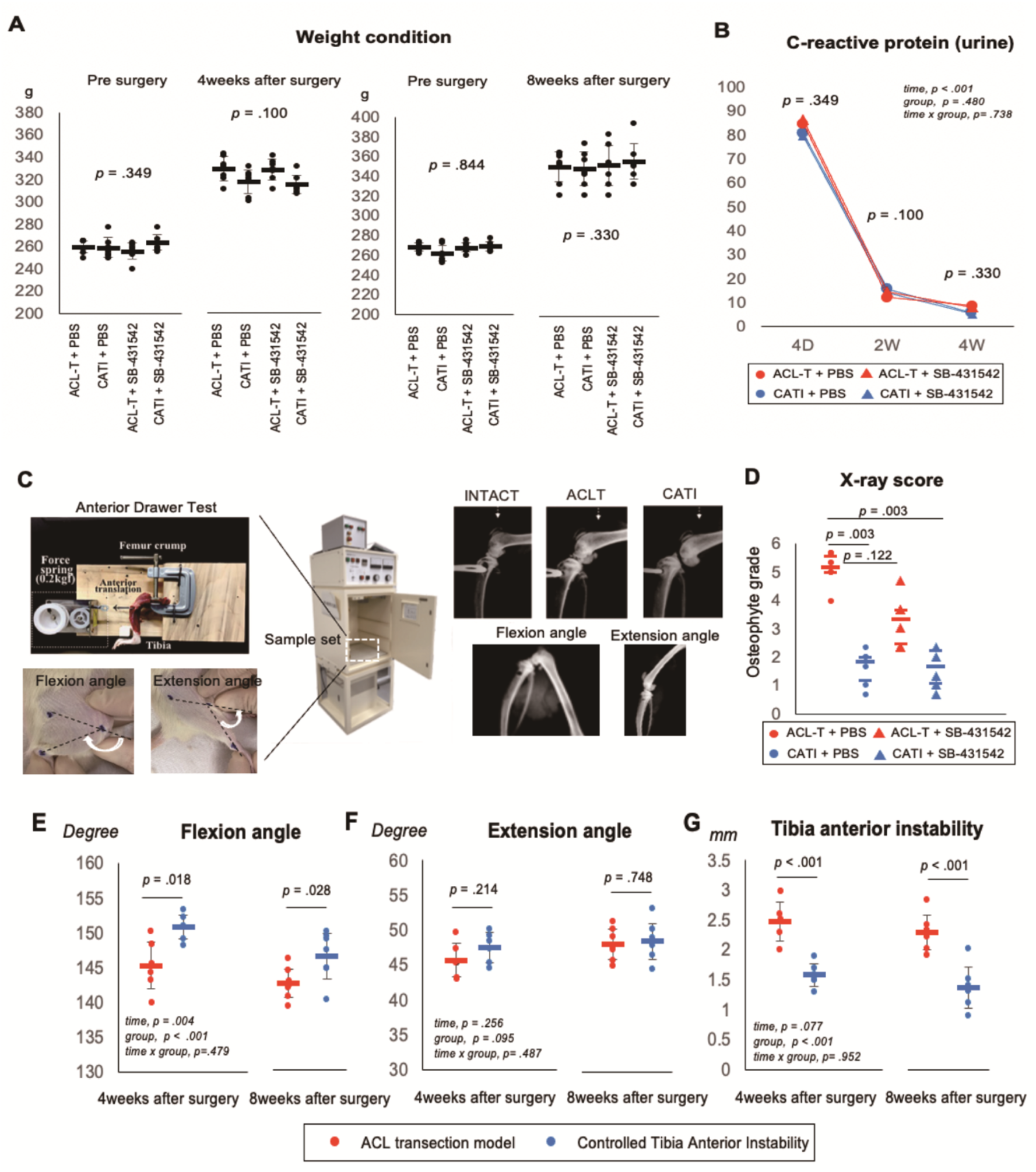
Rat model condition. (A) Body weight changes in each surgical group are shown. There is no significant difference in the effects of surgery and injection in all groups. (b) C-reactive protein (CRP) change shown at 4day, 2 weeks, and 4 weeks. The effect of each surgery was higher on day 4 in all groups, but there was no difference in treatment between groups. In the 2nd and 4th weeks, the CRP level showed a significantly decrease. (c) Tibia anterior instability and Range of motion, osteophytes analysis using soft X-ray. X-ray data were presented as median and 25th/75th percentile (Kruskal-Wallis test, post-hoc Bonferroni correction), weight condition data were presented as means and 95% CI (One-way ANOVA, post-hoc Tukey test), C-reactive protein, angle, and joint instability data also were presented as means and 95% CI (Two-way ANOVA, post-hoc Tukey test)

### 3.2. Biomechanical test

The CATI group showed a slightly restricted flexion range of motion at 4 weeks (ACL-T:145.3 [142.6-148.0] and CATI: 150.8 [149.4-152.3], *p* = 0018; **Fig.2E**) and 8 weeks (ACL-T:142.4 [140.3-144.5] and CATI: 146.3 [143.1-149.5], *p* = 0028; **Fig.2E**). However, there was no difference in the extension ROM in CATI and ACL-T groups a 4 (ACL-T:45.7 [43.3-48.1] and CATI: 47.5 [45.4-49.6], *p* = 0.214; **Fig.2F**) and 8 weeks (ACL-T:48.1 [46.3-49.8] and CATI: 48.5 [46.5-50.5], *p* = 0.748; **Fig.2F**). Regarding the anterior tibia instability, the CATI group was significantly decreased by ACL-T group at 4 (SHAM:45.7 [43.3-48.1] and CATI: 47.5 [45.4-49.6], *p* < 0.001; **Fig.2G**) and 8 weeks (ACL-T:2.5 [2.2-2.7] and CATI: 1.6 [1.4-1.7], *p* < 0.001; **Fig.2G**).

### 3.3. Osteophytes score

Knee Osteophytes were evaluated by two blind evaluators using soft X-ray. The ACL-T group scored higher than the CATI group regardless of the inhibitor treatment (ACL-T:5.0 [5.2-5.6], CATI: 1.8 [1.2-2.0], ACL-T with inhibitor:3.3 [2.5-3.7] and CATI with inhibitor: 1.7 [1.2-2.3], *p* = 0.003; **Fig.2D**). The ACL-T with inhibitor group decreased osteophytes score compare with ACL-T without inhibitor group, although not significantly (*p* = 0.122).

### 3.4. Cartilage degeneration histological score

At 4 and 8 weeks, the OARSI grading scores were significantly high in the ACL-T group compared with the CATI and INTACT knee joint (used opposite lower leg) (4 weeks: ACL-T: 5.0 [4.3-5.3], CATI: 3.0 [2.5-3.6], ACL-T with inhibitor: 3.7 [2.7-3.9], CATI with inhibitor: 3.0 [2.5-3.6] and INTACT: 1.3 [1.0-1.7]. 8 weeks: ACL-T: 8.0 [7.2-8.8], CATI: 4.7 [4.2-4.7], ACL-T with inhibitor: 8.0 [7.3-8.5], CATI with inhibitor: 4.3 [3.3-5.1] and INTACT: 2.0 [2.0-2.3]; **Fig.3A,B**; see ***Supplementary table; Table.S1***). The trend did not change between the with inhibitor and without inhibitor groups. OARSI score was more associated with the degree of joint instability than injection effects (Pearson = 0.732, *p* < 0.001; **Fig. 3E**).

**Fig. 3.**
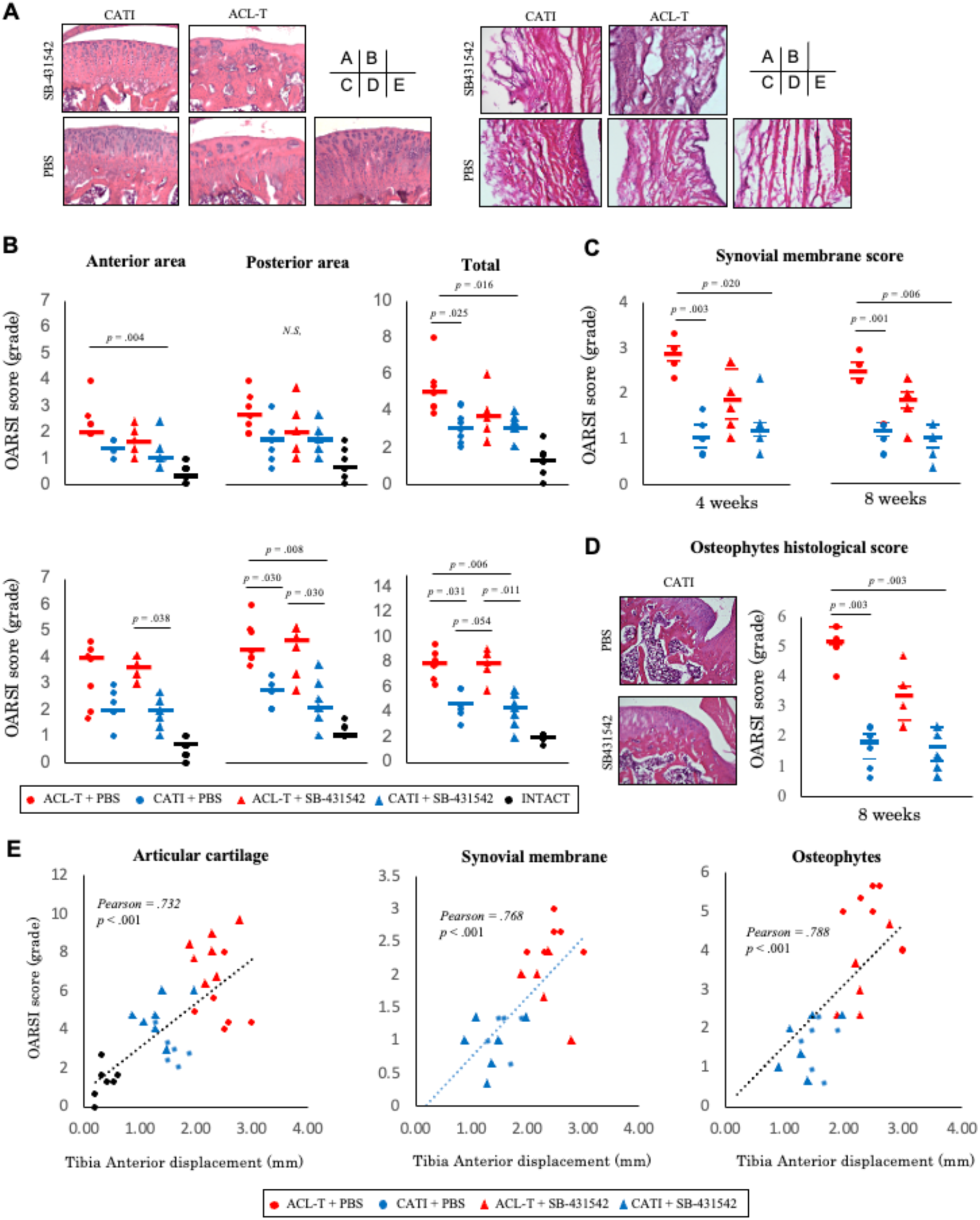
Controlled joint instability inhibit OA pathogenesis in rat. (A) Presented are representative hematoxylin staining images of cartilage (Scale bars: 200 μm) and synovial membrane sections (Scale bars: 100 μm) and (B) quantitation of the OARSI grade scores for histological osteoarthritic findings, (C) synovitis and (D) Osteophytes. Dot plot have shown extending from the 25^th^ (lower bar) and 75^th^ (upper bar) percentiles containing the median. Data used that Kruskal-Wallis, post-hoc Mann-Whitney U test, and Bonferroni correction. (E) Correlation diagram of each histological score and joint instability.

### 3.5. Synovitis histological score

Synovitis scores were suppressed in the both CATI surgery groups with dampened joint instability compared with the ACL-T surgery group. The ACL-T with inhibitor group showed less fibrosis than in the ACL-T without inhibitor group, although the difference was not significant (4 weeks: (ACL-T: 2.8 [2.7-3.0], CATI: 1.0 [0.8-1.3], ACL-T with inhibitor: 1.8 [1.4-2.5] and CATI with inhibitor: 1.2 [1.0-1.3]; 8 weeks: (ACL-T: 2.5 [2.3-2.7], CATI: 1.2 [1.0-1.3], ACL-T with inhibitor: 1.8 [1.7-2.0] and CATI with inhibitor: 1.0 [0.8-1.3]; **Fig.3A,D**). Synovitis score was more associated with the degree of joint instability than inhibitor effects (Pearson = 0.768, *p* < 0.001) (**Fig.3E**).

### 3.6. Osteophyte formation histological score

The osteophyte formation score in the posterior articular cartilage was significantly lower in the CATI than ACL-T surgery groups at 8 weeks, regardless of the injection treatment (ACL-T: 0.15 [0.10-0.21], CATI: 0.42 [0.31-0.53], ACL-T with inhibitor: 0.24 [0.15-0.33] and CATI with inhibitor: 0.35 [0.24-0.45], *p* = .127; Fig.3A,D) (**Fig.3E**). Osteophyte score was also associated with the degree of joint instability than injection effects (Pearson = 0.788, *p* < 0.001) (**Fig. 3E**).

### 3.7. Lubricin expression using IHC

Lubricin (superficial zone protein) associated with articular cartilage lubricity was assessed via IHC evaluation. Both ACL-T surgery groups showed decreased lubricin positive chondrocyte compare with the CATI surgery group in the surface zone (ACL-T: 3.0 [2.0-3.8], CATI: 2.0 [1.0-2.0], ACL-T with inhibitor: 2.0 [1.5-2.8] and CATI with inhibitor: 1.0 [1.0-1.8]; **Fig.4**). No significant score differences were observed between the ACL-T with inhibitor groups and ACL-T without inhibitor. However, the deep area score remained unchanged among all groups (ACL-T: 0.23 [0.09-0.38], CATI: 0.30 [0.20-0.40], ACL-T with inhibitor: 0.29 [0.19-0.39] and CATI with inhibitor: 0.25 [0.18-0.33]; **Fig.4**). Lubricin score was associated with the degree of joint instability than inhibitor effects (Pearson = -0.522, *p* = 0.018).

**Fig. 4.**
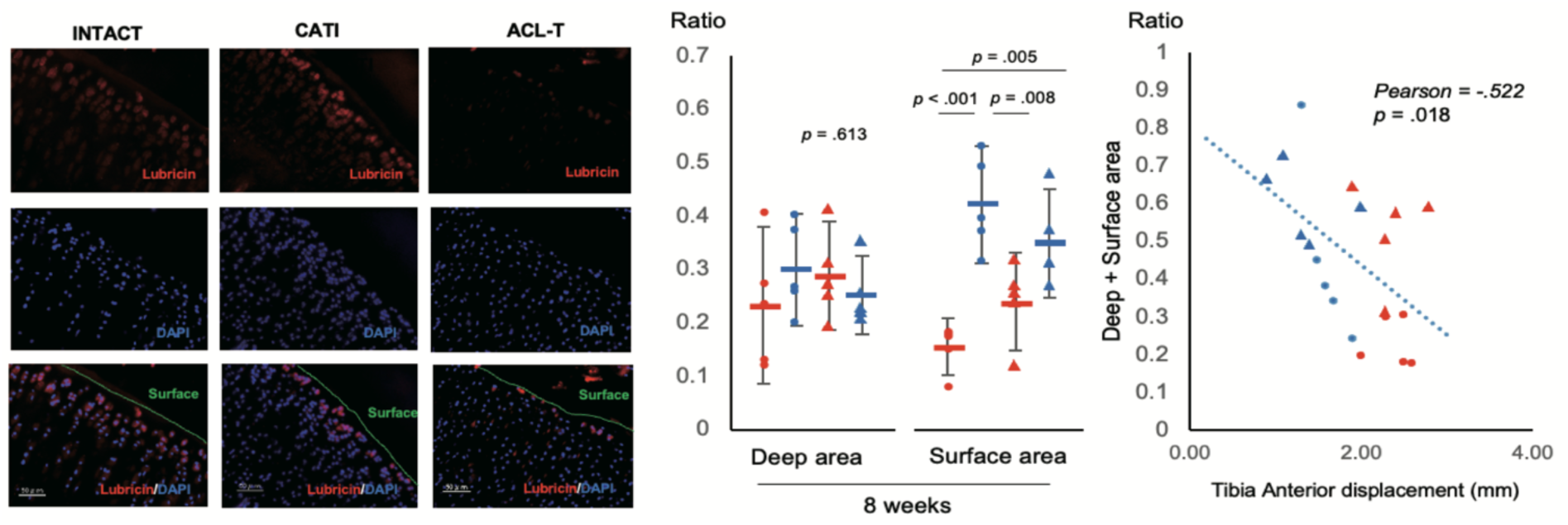
Controlled joint instability maintains cartilage lubricity. IHC staining of lubricin in the articular cartilage and its density are indicated and shown correlation diagram of each histological score and joint instability (Scale bars: 50 μm). To sum up, the joint instability decreased the lubricin in surface cartilage. Data were presented as means and 95% confidence interval (One-way ANOVA, post-hoc Tukey test).

### 3.8. Collagen protein volume expression in synovial membrane

The standardized collagen type I protein concentration was calculated from the measured standard concentration using manufacture instruction. The ACL-T without inhibitor group significantly increased compare with the remaining groups (ACL-T: 5.49 [4.69-6.29], CATI: 3.89 [3.34-4.44], ACL-T with inhibitor: 2.08 [1.56-2.60] and CATI with inhibitor: 2.22 [1.53-2.91]; **Fig. 5A**). Both inhibitor injection groups showed inhibited collagen expression compare with those without the inhibitor treatment. Collagen type Ⅰ expression was associated with the degree of joint instability (Pearson = 0.518, *p* = 0.019; **Fig. 5A**). However, in collagen type Ⅲ, no significant differences were observed in all groups (ACL-T: 34.6 [26.0-43.2], CATI: 25.1 [18.8-31.4], ACL-T with inhibitor: 23.6 [11.6-35.5] and CATI with inhibitor: 15.9 [8.23-23.6], *p* = 0.015 ; **Fig. 5B**); collagen type Ⅲ expression was associated with the degree of joint instability (Pearson = 0.598, *p* = 0.005; **Fig. 5B**). Supplemental figure show the protein expression of IL-1β and TNF-α.

**Fig. 5.**
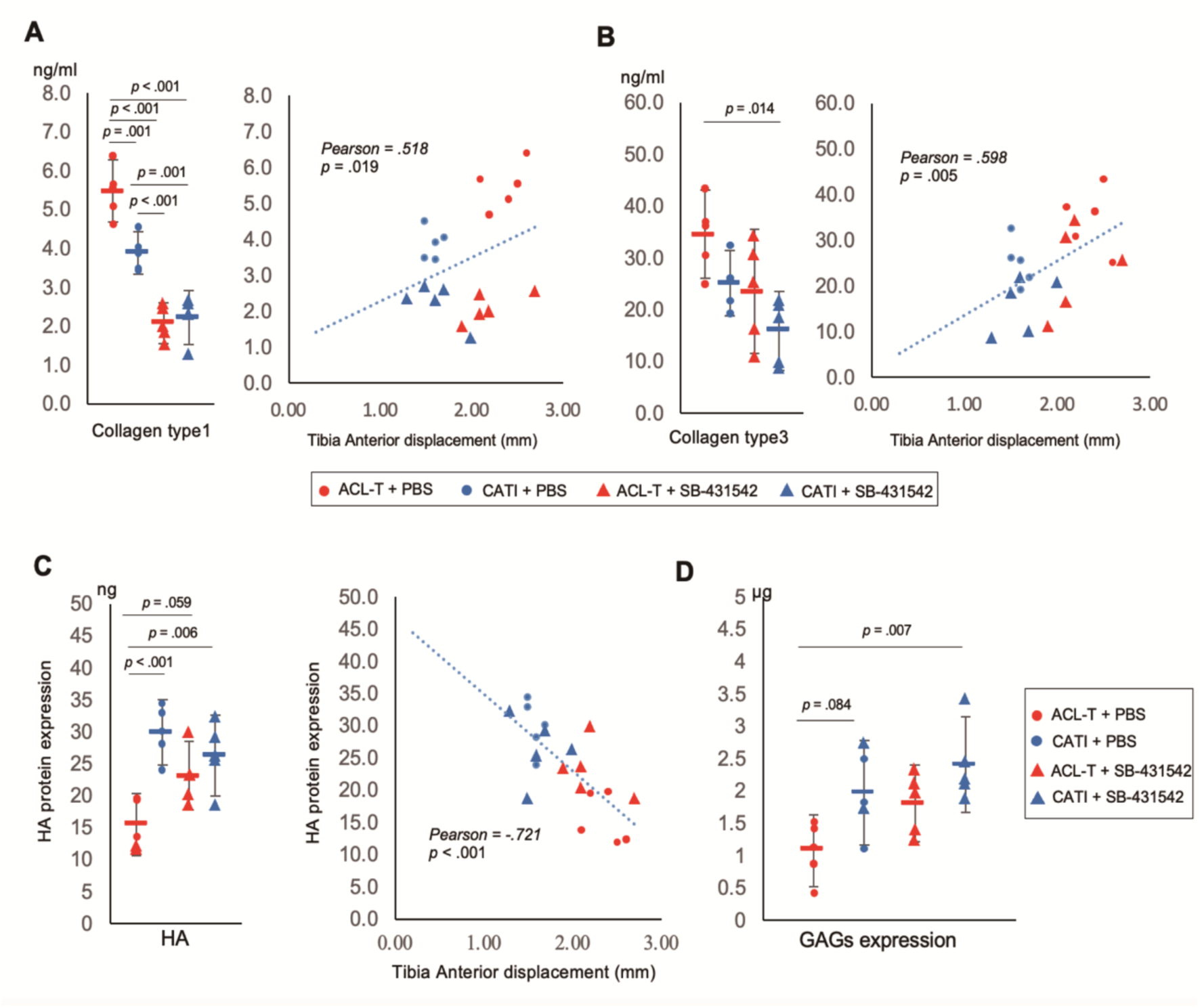
Collagen, HA and GAGs protein volume expression in synovial membrane. (A) Collagen type1 protein expression was determined by ELISA in each experimental synovial membrane tissue (n = 5, in each group. One-way ANOVA, post-hoc Tukey). Moreover, correlation of protein expression ELSA level and joint instability was shown ELISA level. (B) Collagen type3 protein is also determined by ELISA in each experimental synovial membrane tissue (n = 5, in each group. One-way ANOVA, post-hoc Tukey). (C) HA protein expression was determined by ELISA in each experimental synovial membrane tissue (n = 5, in each group. One-way ANOVA, post-hoc Tukey). Moreover, Pearson’s correlations of protein expression ELISA level and joint instability was shown. (D) GAGs protein is also determined by colorimetric method in each experimental synovial membrane tissue (n = 5, in each group. One-way ANOVA, post-hoc Tukey).All data are presented as means and 95% confidence interval.

### 3.9. HA and GAGs protein volume expression in synovial membrane

The standardized HA protein concentration was calculated from the measured standard concentration using the manufacturer’s instruction (B1000, Biocolor Ltd., USA, Assay range: 0.625–50 ng/mL). The HA level in the ACL-T group significantly increased compare with both CATI groups (ACL-T: 15.5 [10.6-20.3], CATI: 29.9 [24.8-35.0], ACL-T with inhibitor: 23.2 [17.8-28.6] and CATI with inhibitor: 26.3 [20.0-32.7]; **Fig. 5C**). Meanwhile, the HA levels in the ACL-T group tended to increase compared with the ACL-T with inhibitor, although not significantly (*p* = 0.059). HA expression was associated with the degree of joint instability (Pearson = -0.721, *p* < 0.001; Fig. 5C). GAGs expression was also evaluated using colorimetric method, which tended to increase in CATI and CATI with inhibitor groups compared with the ACL-T group (ACL-T: 1.08 [0.53-1.63], CATI: 1.98 [1.17-2.79], ACL-T with inhibitor: 1.81 [1.22-2.40] and CATI with inhibitor: 2.41 [1.67-3.15]; **Fig. 5D**).

### 3.10. *mRNA* expression in vitro study

We performed Real-time qPCR to determine whether IL-1β and TGF-β1 can induce synovial origin fibroblast to express collagen and HA synthesis factors (**Fig. 6A**). Addition of IL1β significantly increased the expression level of *IL-6 mRNA* and deceased *PRG4*, *HAS1*, and *HAS2*. However, the results were not affected by TGF-β1, therefore, synovial origin fibroblast is regulated independently of IL1-β and TGF-β1. The TGF-β1 inhibitor, SB-43152, significantly decreased collagen, however, had weak effects on other factors (see ***Supplementary table; Table.S2 and S3***).

**Fig. 6.**
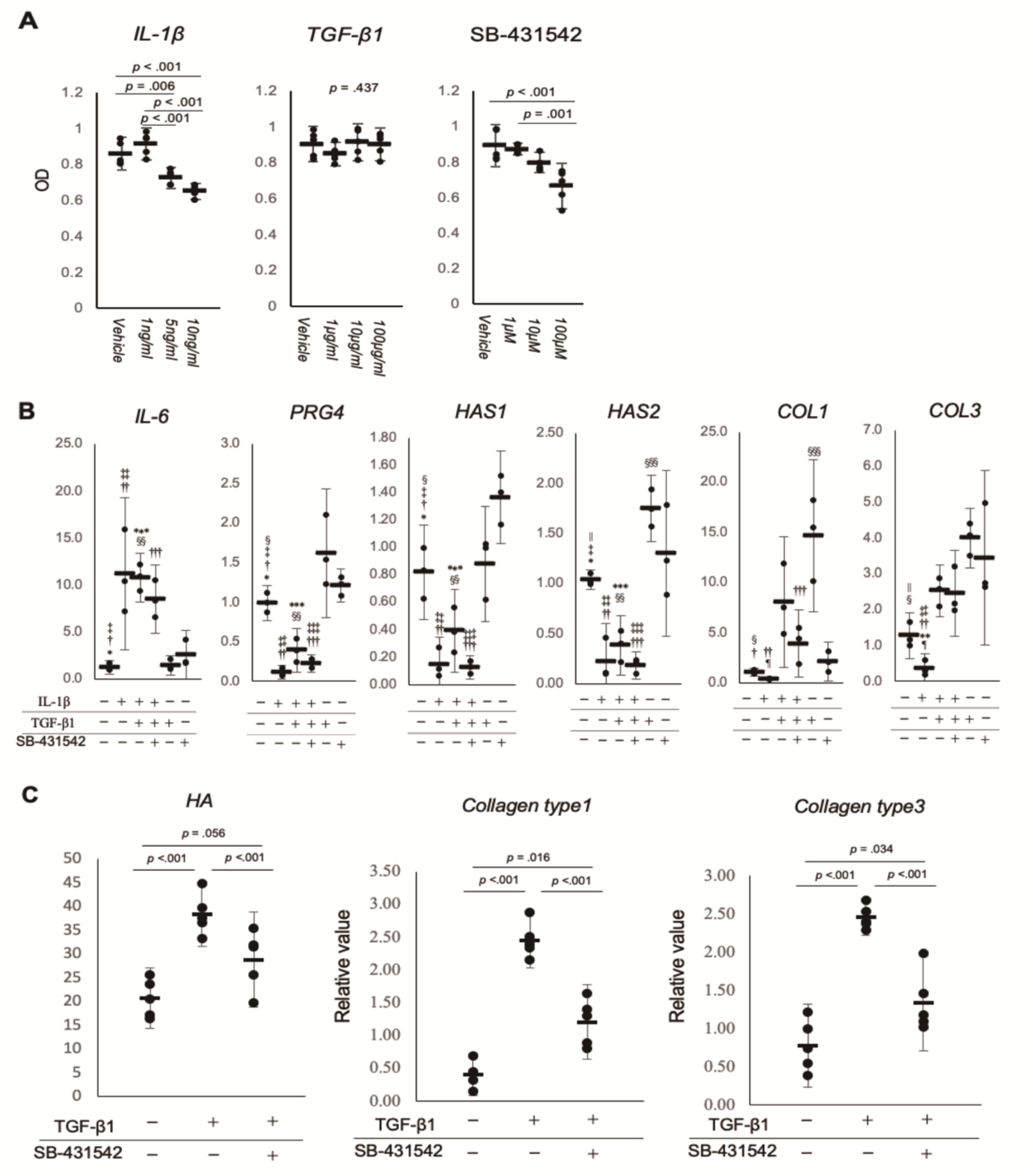
Inhabitation of TGF-β1 decreased fibrosis factors and HA on fibroblast. (A) Cell proliferation ability in each treatment using CCK-8 (n = 5, in each group. One-way ANOVA, post-hoc Tukey). (B) Fibroblast after treatment mRNA expression. The figure showed relative value using ΔΔCt methods. Statistical analysis was performed using delta Ct values using GAPDH and actin housekeeping (n = 3, in each group. One-way ANOVA, post-hoc Tukey). Analyses were performed at 24 h. *, †, ‡, §, ||, ¶, **, ††, ‡‡, §§, ***, †††, ‡‡‡= *p* < 0.05 and indicates significantly different from other groups. Detail p-value is shown Supplementary table; Table.S2 and S3. (C) ELISA results for the expression of protein in the synovial membrane about to analysis role TGF-β. Data are presented as means (black bar) and 95% confidence interval (n = 5, in each group. One-way ANOVA, post-hoc Tukey).

### 3.11. Collagen analysis by TGF-β stimulation in fibroblasts

The protein levels of collagen type Ⅰ, III and HA were investigated using synovial fibroblasts. The medium containing TGF-β had increased collagen types Ⅰ and Ⅲ and HA compare with those no TGF-β1 (HA: Vehicle, 20.7 [9.7-22.6]; TGF-β1, 38.4 [11.2-24.9]; TGF-β1+SB-431542, 28.8 [28.9-48.9]. Collagen type1: Vehicle, 0.40 [0.08-0.72]; TGF-β1, 2.45 [2.02-2.88]; TGF-β1+SB-431542, 1.20 [0.64-1.77]. Collagen type3: Vehicle, 0.77 [0.24-1.31]; TGF-β1, 2.45 [2.21-2.89]; TGF-β1+SB-431542, 1.34 [0.71-1.97]; **Fig. 6C**). The medium containing TGF-β1 with TGF-β1 inhibitor inhibited both collagen and HA synthesis (HA: *p* = 0.056, collagen type Ⅰ: *p* = 0.016, collagen type Ⅲ: *p* = 0.034).

## 4. Discussion

This study aimed 1) to determine whether cartilage degeneration is inhibited in our model in which joint instability is controlled, 2) to determine whether TGF-β1 inhibition mitigates cartilage degeneration, and 3) to determine the role of TGF-β1 in synovitis using fibroblasts from the synovial membrane. Our results showed the negative role of joint instability in knee joint tissue such cartilage and osteophyte, and that the over expression of TGF-β1 in synovitis increased collagen and fibrosis.

Joint instability is a risk factor for OA onset and progression. In patients with OA, knee instability is associated with significant alterations in knee joint dysfunction, and induced joint dysfunction during gait [39,40]. Moreover, knee joint instability compromises function in individuals with medial knee osteoarthritis and may be related to impaired joint mechanics [41]. Conversely, in animal models, joint instability induced by knee ligament alters kinematics and OA pathology. Kamekura et al. have reported that ACL, MCL, LCL, and Meniscus transected in different combinations cause various degrees of joint instability, which is associated degeneration [42]. ACL transection (ACL-T) model also causes joint instability, thereby promoting cartilage degeneration [43]. However, previous studies have failed to demonstrate that joint instability is an independent cartilage degeneration factor, since various ligaments were transected. To address this gap, our CATI model provided external stability in the ACL-T model, while joint instability by ACL injury was controlled. In this study, we observed a clear difference in the articular cartilage degeneration and osteophytes formation between ACL-T and CATI in at 8 weeks. Moreover, synovial membrane in the CATI group inhibited inflammatory cytokines compare with the ACL-T group.

The major changes in the histological analysis and synovial protein levels between the two models indicate that joint instability independently controls joint conditioning. Biomechanical change at the knee joint is a complex phenomenon, knee stability is attributed to a combination of static structures and biomechanical changes in daily living. Knee ligament prevents instability, while ACL injury is a main stabilizer of the knee. Joint instability due to ACL injury induced biomechanical changes; our model limits abnormal joint instability. To sum up, our results indicate that joint instability promotes synovitis and cartilage degeneration.

To evaluate role of TGF-β1 cascade in knee joint conditions, we used the inhibitor SB-431542. Although the lubrication of synovial fluid produced by HA are essential to normal movement of the knee joint, HA synthesis in the synovial membrane is affected by various mediators. For example, TGF-β1 expression in the knee synovial membrane positively influences HA synthesis in the synovium, hence maintains lubrication in article cartilage. The over expression of TGF-β1 affects collagen expression through the activation of the synovial cells and thickening of the synovial tissue. Chronic synovial inflammation in OA may induce structural degeneration progression, hence improving our knowledge of synovial physiology is critical to designing safe and effective OA treatments. Our study investigated the influence inhibiting the TGF-β1 pathway in knee joints using micro injection on cartilage and synovium. Although the inflammatory factors in the synovial membrane increased following ACL-T surgery, SB-431542 injection inhibited pro-inflammatory factors such as TNF-α and IL-1β. Moreover, HA and GAGs protein expression increased by inhibitor injection compared to that by ACL-T surgery. The synthesis of collagen type Ⅰ is controlled by the TGF-β1 protein pathway. A previous study on synovial fibroblasts also showed that TGF-β1 signaling is an inducer of synovial tissue fibrosis, and proliferates types Ⅰ and 3 collagen, and that synovial macrophage induced osteophyte formation through synovium mediating pathway [44]. Our study indicated that joint instability induced collagen synthesis in synovial membrane associated with synovitis and osteophyte, and TGF-β1 inhibitor decreased collagen synthesis factor, with maintenance of HA and GAGs. These results suggest that suppressing synovitis caused by joint instability can suppress synovial fibrosis, osteophytes, and degeneration of cartilage.

Whether changes in TGF-β1 directly affect the cartilage and synovium remains unclear. Therefore, as assessed fibroblast from synovial membrane in inflammation condition, to determine the role of TGF-β1. IL1β alone did not influence the TGF-β1 effect on fibroblasts, which is similar to previous findings on skin and lung fibroblasts exposed to TGF-β1 [45]. Meanwhile, addition of TGF-β1 to fibroblast increased collagen mRNA and protein. Moreover, the TGF-β1 signaling pathway involves the phosphorylation of Smad2 and Smad3 proteins through activin receptor-like kinase (ALK). Zhu et al. have reported that TGF-β1 stimulated type Ⅱ collagen and aggrecan expressions in rat chondrocytes, while treatment with ALK5 inhibitor significantly impaired TGF-β1-induced type Ⅱ collagen and aggrecan expressions [46]. Our results in the synovial membrane showed that TGF-β1 increased HA and GAGs expressions, which were reduced by ALK5 suppression by the SB-431542 inhibitor. These results suggest that joint instability is an independent mechanical factor for OA progression and provide novel insights into the association between OA and joint instability, which has significant clinical implications.

## 5. Conclusions

The present study showed that joint instability exacerbates articular cartilage degeneration and decreases HA and GAGs protein expressions in synovial membrane (**Fig.7**). Stable joint is pivotal in synovial membrane and cartilage stability. Moreover, synovitis induced by joint instability can be inhibited using TGF-β1 inhibitor. Our results also showed that the effects of TGF-β1 inhibitor on joints may suppress synovial inflammation, however, mechanical stress, including joint instability, must also be considered at the same time.

**Fig. 7.**
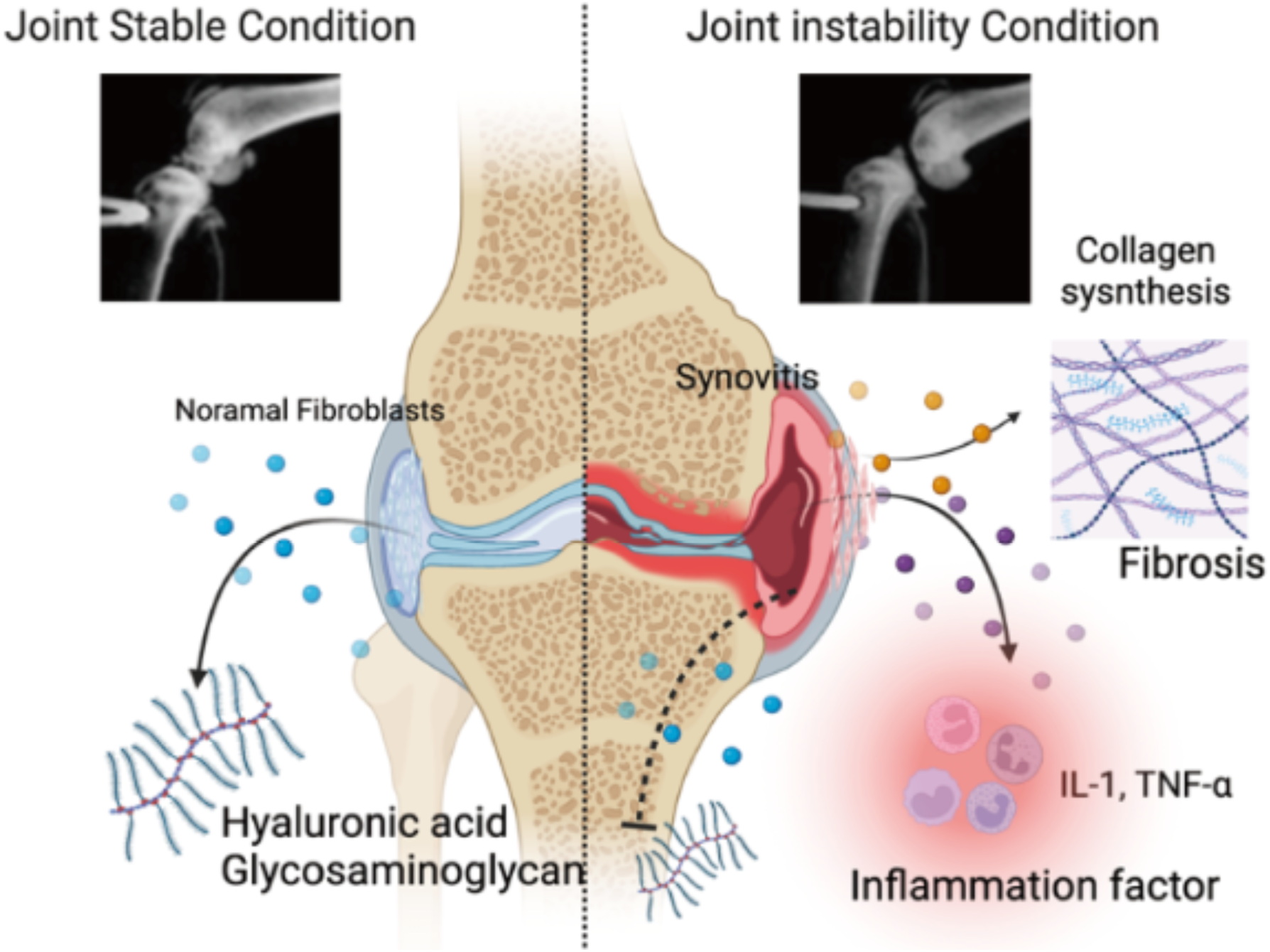
Schema of Joint Instability and Osteoarthritis.

### Compliance with ethics requirements

All Institutional and National Guidelines for the care and use of animals were followed.

## Acknowledgements

This study was supported by JSPS KAKENHI (20K19417) Grant-in-Aid for Young Scientists. Moreover, schematic diagram illustrating the osteoarthritis mechanism associated joint instability (graphic abstract) was generated using Biorender.

## Authorship contribution statement

The authors would like to thank all authors participated in the experiment according to the below, and Asterisks is the main person responsible for each experiment. Finally, all authors approved the final version to be published. Concept and design of the studies composed K.M.*, S.K. and N.K.. In vivo study is mainly carried out K.M. * and followed by T.K, Y.O., and C.T.. They are contributed for blind data in histological analysis and osteophytes analysis. In vitro study is mainly carried out K.M.*, H.T. and T.K.. Drafting the article is K.M., funding acquisition was K.M..

## Supplementary Information

The supplementary tables (S1, S2 and S3) available at https://XXX

The supplementary results (SR) available at https://XXX.

---

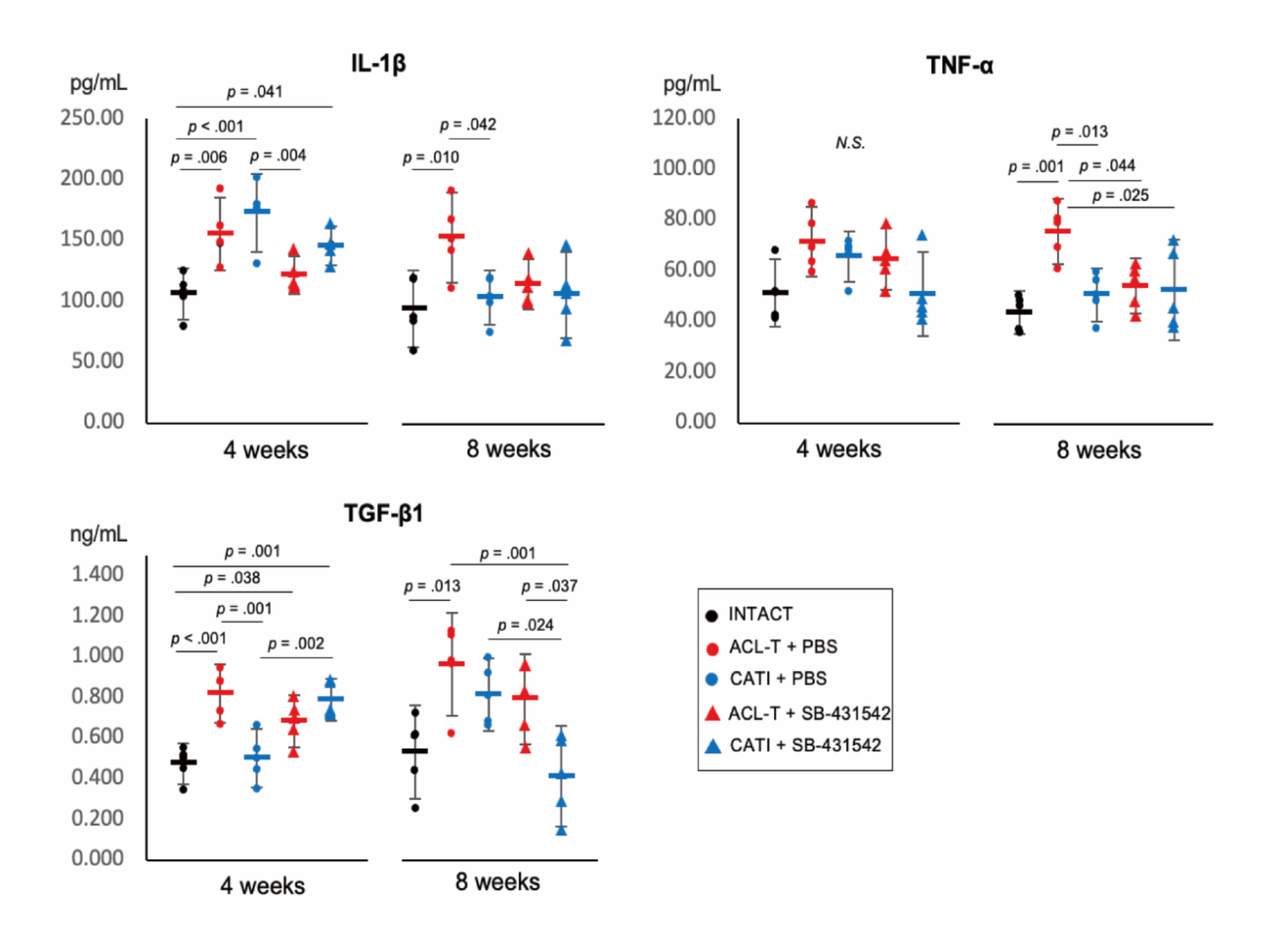
Supplementary figure.

**Supplementary figure information** Supplemental data shows the protein expression values obtained from the synovial membrane samples. Difference of protein expression associate inflammation between CATI and ACL-T groups was no significant IL-1β, and TNF-α. However, protein volume at 8weeks is significant increased ACL-T groups compare than CATI groups. There results might be indicated that joint instability condition associate to continue chronic inflammation in knee. On the other hand, HA expression that maintain factor of articular cartilage is significantly increased CATI group than ACL-T groups. About effect of TGF-β inhibitor that associated fibrosis for synovial membrane was not significant change. TGF-β protein expression is that ACL-T group was significantly highest among all group regardless of injection at 4 weeks. However, the ACL-T+ inhibitor group significantly decreased the expression of type I and type 3 collagen compared to ACL-T. This intriguing result suggests that suppression of TGF into joints may prevent fibrosis.

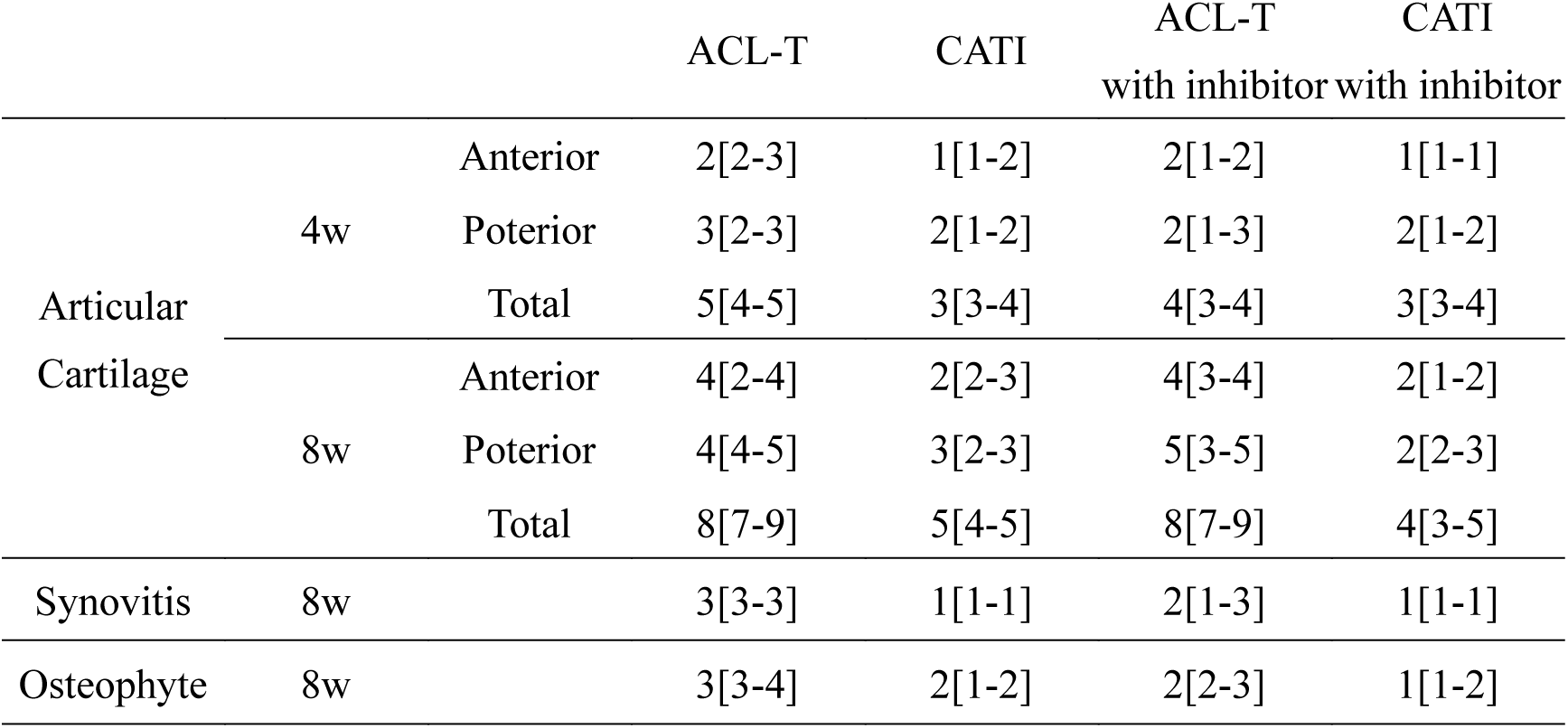
Supplementary *table 1*.

***Supplementary table 1***
|  |  |  | ACL-T | CATI | ACL-T<br>with inhibitor | CATI<br>with inhibitor |
| --- | --- | --- | --- | --- | --- | --- |
| Articular<br>Cartilage | 4w | Anterior | 2[2-3] | 1[1-2] | 2[1-2] | 1[1-1] |
|  |  | Posterior | 3[2-3] | 2[1-2] | 2[1-3] | 2[1-2] |
|  |  | Total | 5[4-5] | 3[3-4] | 4[3-4] | 3[3-4] |
|  | 8w | Anterior | 4[2-4] | 2[2-3] | 4[3-4] | 2[1-2] |
|  |  | Posterior | 4[4-5] | 3[2-3] | 5[3-5] | 2[2-3] |
|  |  | Total | 8[7-9] | 5[4-5] | 8[7-9] | 4[3-5] |
| Synovitis | 8w |  | 3[3-3] | 1[1-1] | 2[1-3] | 1[1-1] |
| Osteophyte | 8w |  | 3[3-4] | 2[1-2] | 2[2-3] | 1[1-2] |

Supplementary *table 2*.

**Supplementary table 2**
|  |  |  |  |  |  |  |  |
| --- | --- | --- | --- | --- | --- | --- | --- |
| | IL-1β | - | + | + | + | - | - |
| | TGF-β1 | - | - | + | + | + | - |
|  | SB-431542 | - | - | - | + | - | + |
|  | *IL-6* | 1.27[0.51-2.03] | 11.20[3.13-19.26] | 10.82[8.26-13.39] | 8.52[4.88-12.15] | 1.47[0.48-2.47] | 2.63[0.03-5.22] |
|  | *prg4* | 1.00[0.78-1.22] | 0.11[0.03-0.20] | 0.40[0.12-0.68] | 0.23[0.12-0.34] | 1.62[0.81-2.44] | 1.21[1.00-1.42] |
|  | *HAS1* | 1.04[0.94-1.14] | 0.22[-0.15-0.60] | 0.38[0.09-0.68] | 0.19[0.05-0.32] | 1.75[1.42-2.09] | 1.30[0.47-2.14] |
|  | *HAS2* | 0.82[0.48-1.16] | 0.15[0.04-0.34] | 0.39[0.09-0.70] | 0.13[0.04-0.21] | 0.88[0.46-1.30] | 1.37[1.03-1.70] |
|  | *COL1* | 1.09[0.70-1.48] | 0.41[0.31-0.50] | 8.08[1.56-14.60] | 3.91[0.57-7.25] | 14.63[7.06-22.20] | 2.12[0.19-4.05] |
|  | *COL3* | 1.28[0.64-1.92] | 0.36[0.04-0.76] | 2.53[1.79-3.26] | 2.45[1.26-3.65] | 4.00[3.17-4.84] | 3.45[1.02-5.88] |
|  |  |  |  |  |  |  | mean [95%CI] |

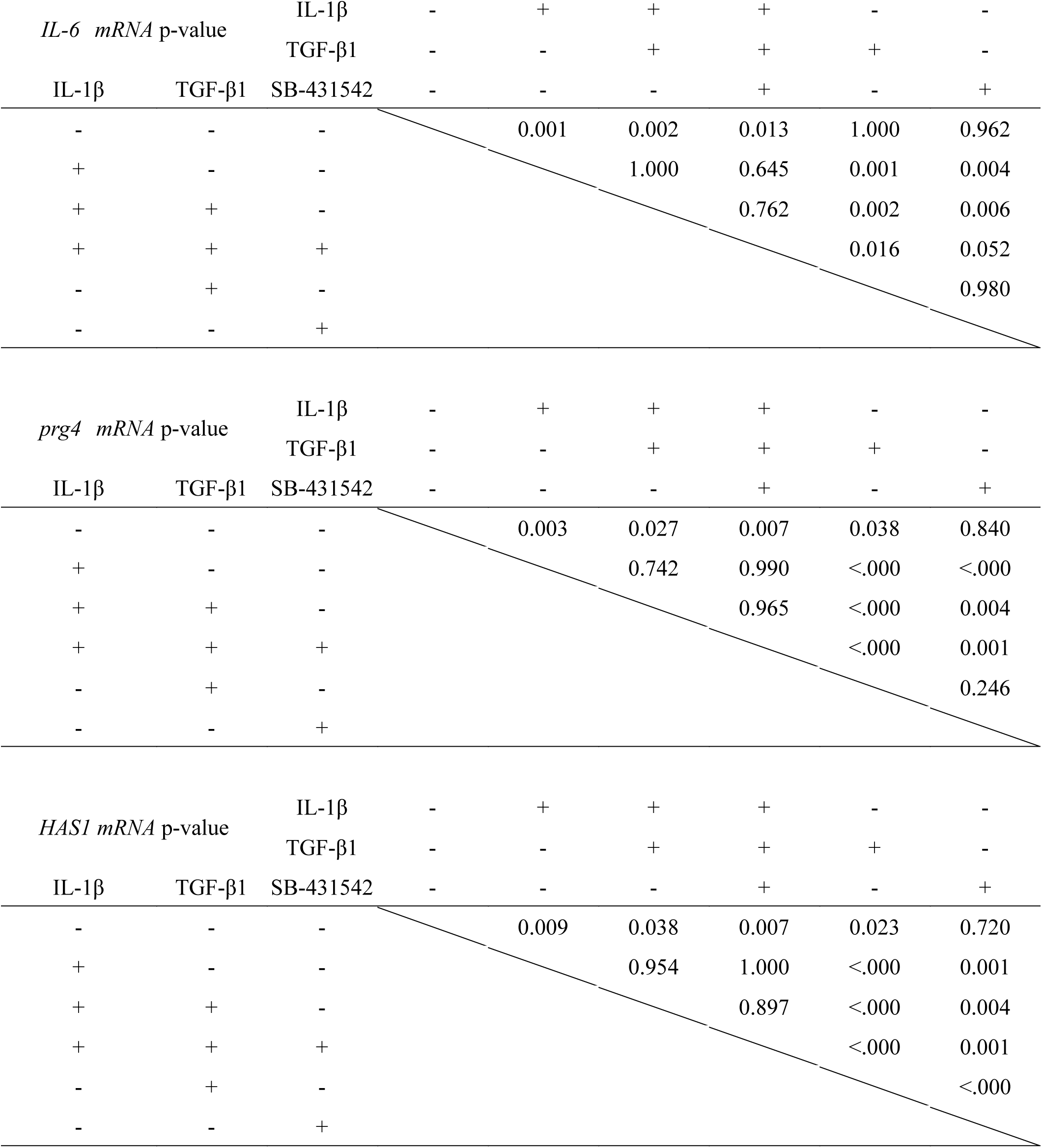
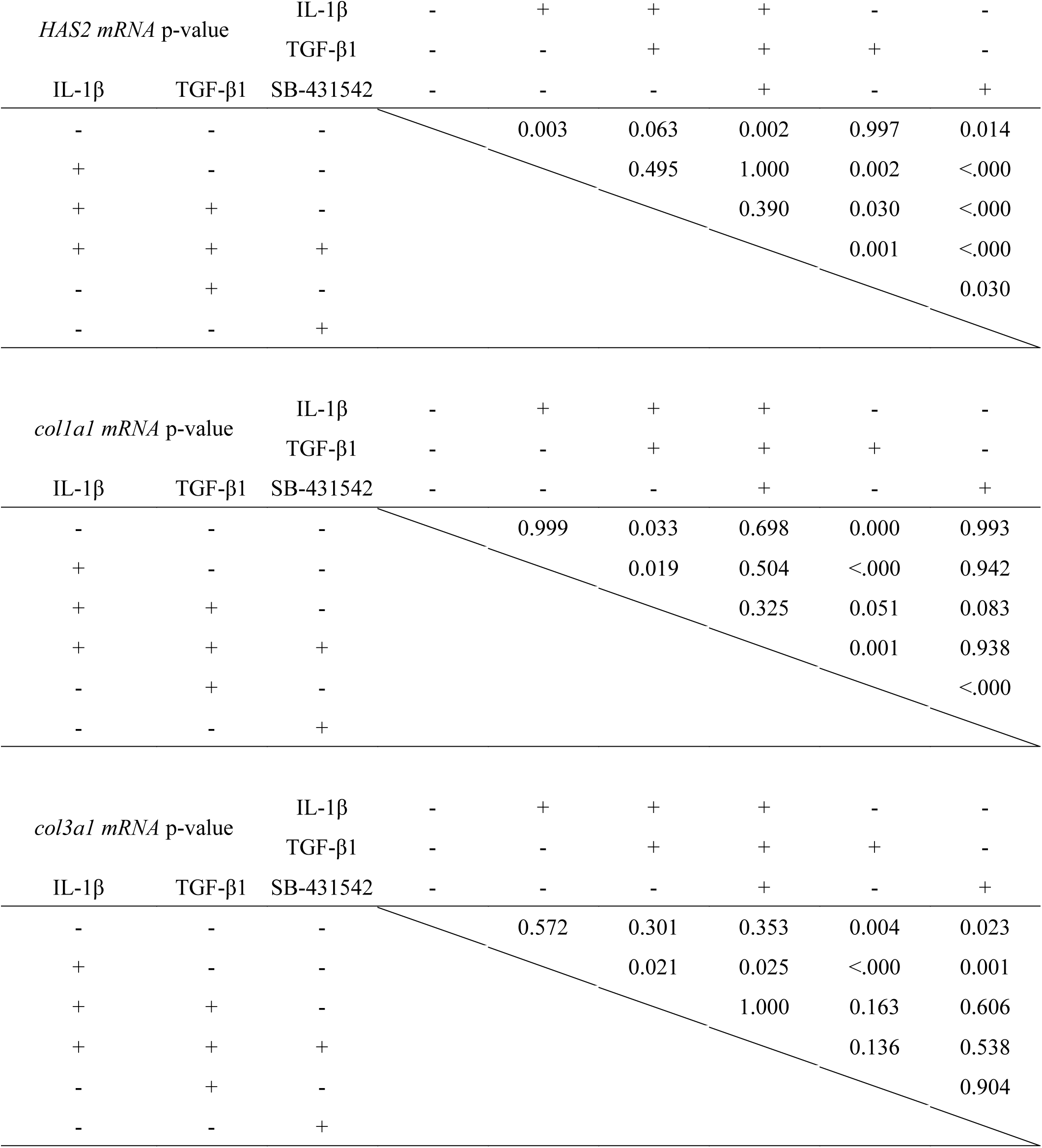
Supplementary *table 3*.

